# The transporters SLC35A1 and SLC30A1 play opposite roles in cell survival upon VSV virus infection

**DOI:** 10.1101/573253

**Authors:** Anna Moskovskich, Ulrich Goldmann, Felix Kartnig, Sabrina Lindinger, Justyna Konecka, Giuseppe Fiume, Enrico Girardi, Giulio Superti-Furga

**Author notes:** Correspondence and requests for materials should be addressed to: Enrico Girardi and Giulio Superti-Furga, CeMM Research Center for Molecular Medicine of the Austrian Academy of Sciences, Lazarettgasse 14, AKH BT25.3, 1090 Vienna, Austria.

## Abstract

Host factor requirements for different classes of viruses are still to be fully unraveled. Replication of the viral genome and synthesis of viral proteins inside the human host cell are associated with altered, often enhanced, cellular metabolism and increased demand for nutrients as well as specific metabolites. With more than 400 members listed to date in humans, solute carriers (SLCs) represent the largest family of transmembrane proteins dedicated to the transport of ions and small molecules such as amino acids, sugars and nucleotides. Consistent with their impact on cellular metabolism, several SLCs have been implicated as host factors affecting the viral life cycle and the cell response to infection. In this study, we aimed at characterizing the role of host SLCs in cell survival upon viral infection by performing unbiased genetic screens using a focused CRISPR knockout library. Genetic screens with the cytolytic vesicular stomatitis virus (VSV) showed that loss of two SLCs genes, encoding the sialic acid transporter SLC35A1/CST and the zinc transporter SLC30A1/ZnT1, affected cell survival upon infection. Further characterization of these genes pointed to a role of both transporters in the apoptotic response induced by VSV, offering new insights into the cellular response to oncolytic virus infections.

## Introduction

Solute carriers (SLCs) comprise the largest family of transporters in the human genome and the second largest family of transmembrane proteins after G-protein coupled receptors (GPCRs). As such, they represent the major “gatekeepers” of energy supply and metabolism in a cell, controlling the uptake and release of a growing list of small molecules and metabolites^1^. With more than 400 members, SLCs are a significantly understudied family considering that ∼30% of family members still have completely unknown function(s) or substrate specificity, despite a large number of links between SLCs and human diseases^1,2^.

Viruses represent a major class of pathogens affecting human health and act as potent and pleiotropic perturbations of cellular metabolism and function, offering the possibility to explore the biology of cellular and immunological processes. Given their complete dependence on host metabolism for replication, viruses have been shown to dramatically alter and exploit cellular pathways for their own purpose^3^. Accordingly, transporters have been shown to be modulated during virus infection^4^, are required as viral entry receptors^5,6^ or as factors affecting viral cell entry^7,8^. Finally, solute carriers can play important roles in determining the outcome of anti-viral immune responses^9,10^.

In this study, we used an unbiased genetic screening approach to identify transporters affecting infectivity and cellular survival upon treatment with influenza virus (IAV) or vesicular stomatitis virus (VSV). In particular, we focus on VSV, which has been extensively used as a model system for virus replication and viral life cycle studies^11^. A cytolytic virus, VSV is able to induce rapidly and effectively cell death. Apoptosis is induced either via the mitochondrial intrinsic pathway via caspase-9 activation, via death receptor-mediated (extrinsic pathway) apoptosis or via ER-stress mediated apoptosis^12^. Due to its non-pathogenicity in humans and its ability to rapidly induce apoptosis in infected cancer cells, VSV has been showing promise in cancer immunotherapies^13^ and oncolytic virotherapies^12^, as well as for the development of vaccines e.g. against Ebola virus^14^. Therefore, there is significant interest in determining the host factors affecting cell survival upon VSV infection. Here we describe two transporters, the sialic acid transporter SLC35A1 and the zinc transporter SLC30A1 (ZnT1), as factors promoting resistance or sensitivity upon VSV infection, respectively.

## Results

### A SLC-focused CRISPR knockout genetic screen identifies SLC35A1 and SLC35A2 as factors required for IAV infection

In order to identify solute carriers affecting viral infection and cell survival, we performed unbiased genetic screens using a SLC-focused CRISPR knockout (KO) lentiviral library targeting 388 SLC genes with multiple sgRNAs per gene^15^ (Fig 1a). To test the ability of this library to identify factors affecting viral infection, we screened for transporters affecting survival of the lung adenocarcinoma cell line A549 upon influenza A virus (IAV, strain A/WSN/33) infection. A549 cells were transduced with the SLC KO library and subsequently infected with IAV at the MOI of 0.5. Samples were collected 96 hours post infection (h.p.i.) and the library compositions of the treated versus untreated samples were compared after recovery and sequencing of sgRNA inserts. We observed good representation of the library in all samples (Suppl Fig 1a), and good correlations between replicates (Suppl Fig 1b). Consistent with previous results^16,17^, we identified two SLC genes, *SLC35A1* and *SLC35A2*, whose sgRNAs were significantly enriched in the surviving cell population (Fig 1b, Suppl Table 1). Both genes are members of the large SLC35 family of nucleotide-sugar transporters and the two corresponding proteins localize to the Golgi apparatus^18^. SLC35A1/CST is a CMP-sialic acid/CMP antiporter while SLC35A2 is a UDP-galactose/UDP antiporter. IAV exploits sialic acid, a hexose residue present on several oligosaccharides, as its main cell entry receptor^19^. These transporters have been previously shown to play an important role in determining the glycosylation pattern of secreted and plasma membrane proteins and are associated with congenital disorders of glycosylation^20,21^. In particular, lack of SLC35A1 or SLC35A2 (which results in the loss of the galactose residue often present upstream of the terminal sialic acid) has been shown to cause reduced or abolished levels of sialyation on the cell surface, resulting in a severe impairment of IAV docking and entry^16,17^. Consistent with the screening results, CRISPR/Cas9-based knockout of the *SLC35A1* or *SLC35A2* genes in A549 cells resulted in increased resistance to IAV infection (Fig 1c, Suppl Fig1c-d). The *SLC35A1* inactivation effect could be reversed by ectopic expression of SLC35A1 (Fig 1c, Suppl. Fig 1e), confirming the dependence of the observed phenotype on this specific gene. Altogether, this set of experiments validated the use of our SLC-focused CRISPR knockout library as well as our experimental approach to investigate the role of SLC genes in the viral life cycle in tissue-cultured cells. Moreover, it confirmed the non-redundant role of members A1 and A2 of the SLC35 family, among all other SLCs represented in the library, on determining the ability of influenza A viruses to infect human cells.

**Fig. 1.**
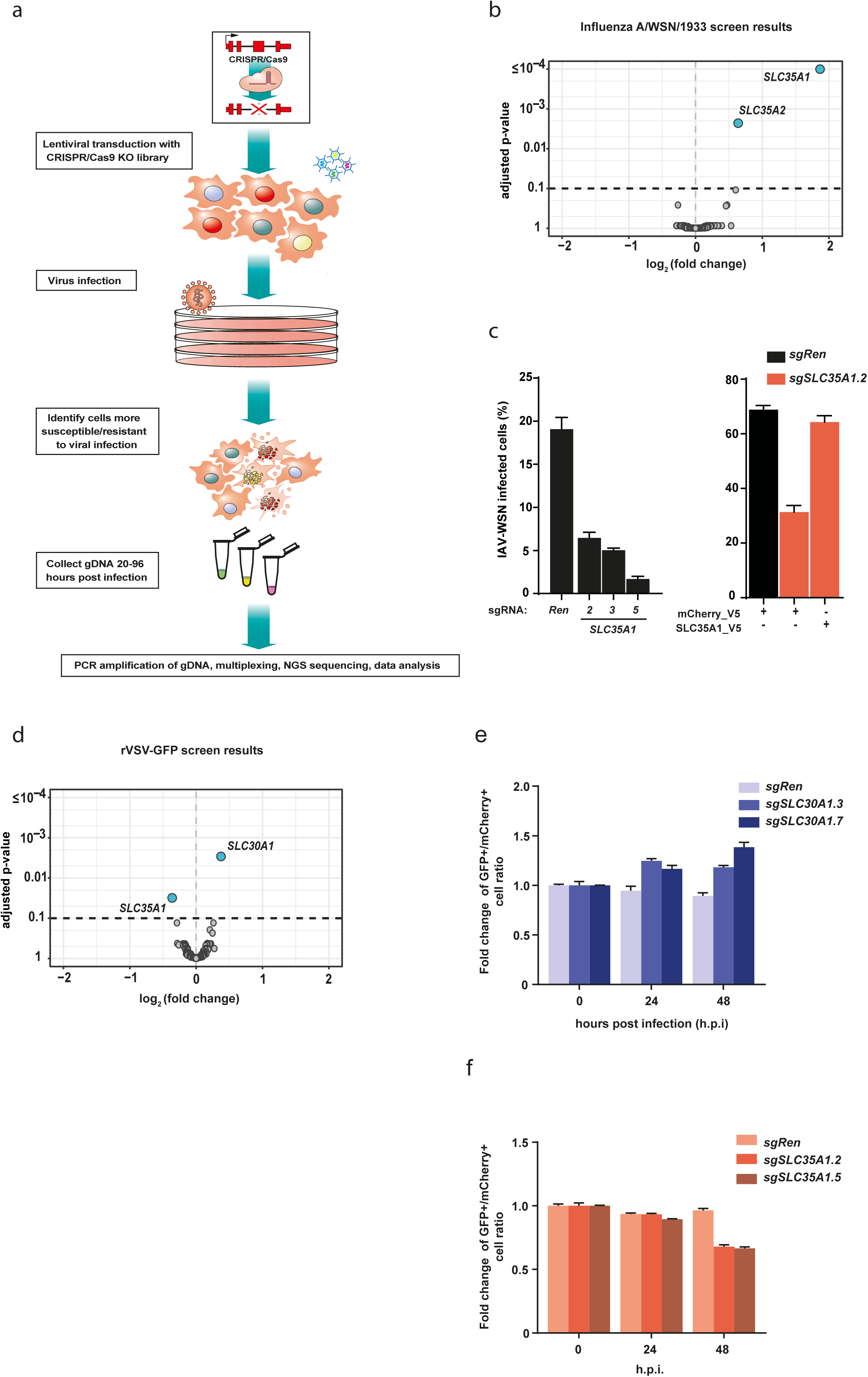
A SLC-focused genetic screen identifies known and novel host factors for IAV and VSV. **a**. Schematic overview of the screen workflow. **b.** Volcano plot of the genetic screen results upon infection of A549 cells with IAV-WSN/1933 at 96 h.p.i. versus uninfected cells **c.** Quantification of flow cytometry-based viral replication assay for selected hits. A549 cells carrying sgRNAs targeting *SLC35A1* or *Renilla* luciferase control as well as cell expressing SLC35A1 cDNA were infected with IAV-WSN/33 at MOI 0,01 and 24 h.p.i. the number of the infected cells was assessed after staining with anti-Influenza Nuclear protein antibody coupled to AF488. **d.** Volcano plot of the genetic screens results in A549 cells infected with VSV-GFP at 48 h.p.i. **e-f.** Multicolour competition assay (MCA) of *SCL30A1* (**e.**) or *SLC35A1* (**f.**) knock-out cells and *Renilla* controls. Cells were mixed at 1:1 ratio, infected with VSV-rWT at MOI 5 and cultured until the indicated time points. The percentage of mCherry+ (*Renilla* control) and eGFP+ (gene of interest) cells in the live population (FSC/SSC) were assessed using flow cytometry. Representative results from at least two independent experiments performed in triplicates.

### Genetic screening identifies SLC35A1 and SLC30A1 as modulators of cell survival upon VSV infection

A549 cells showed substantial cell death 24 hours post VSV infection (above 50% at MOI 10) (Suppl Fig 1f). For the genetic screen, A549 cells were infected with the SLC library and subsequently infected with VSV virus at a MOI of 10 (Fig 1a). Samples were collected pre-infection and at 48 h.p.i. As before, we observed good representation of the library in all samples (Suppl Fig 1g), and good correlations between replicates (Suppl Fig 1h). Comparison of the sgRNA abundance between the virus-treated and control samples at 48h showed a moderate but significant differential enrichment for two transporters: *SLC30A1* (ZnT1, log_2_ fold change: 0.373, adjusted *p*-value: 7×10^-6^) and *SLC35A1* (log_2_ fold change: - 0.362, adjusted *p*-value: 1.5×10^-4^) (Fig 1d). Interestingly loss of SLC35A1, which conferred resistance in the IAV screen, resulted in increased cell sensitivity upon VSV infection. Moreover, we identified as a permissive factor SLC30A1, the only member of the SLC30 family reported to export zinc at the plasma membrane^22^. In order to validate these results, we used a multicolor competition assay (MCA) approach in A549 cells carrying sgRNAs targeting *SLC30A1, SLC35A1* or expressing a sgRNA targeting *Renilla luciferase* as negative control. We observed a modest but clear enrichment of cells carrying *SLC30A1* mutations compared to control cells at both 24 and 48 hours after infection, validating the outcome of the screen and suggesting that indeed SLC30A1 is exerting a negative effect on cell survival after VSV infection (Fig 1e, Suppl Fig 1i). Conversely, inactivation of the *SLC35A1* gene resulted in increased cell death compared to control at 48 hours, suggesting that this transporter support cell survival after VSV infection thereby validating the primary screen (Fig 1f, Suppl Fig 1j).

### Loss of the Zn-exporter SLC30A1 inhibits caspase activation upon VSV virus infection

SLC30A1 has been reported to be responsible for the export of zinc from the cytoplasm^22,23^. To confirm its role in the homeostasis of zinc levels in our cellular system, we measured intracellular levels of zinc using the Zinpyr1 dye^24^. A549 cells lacking SLC30A1 (Suppl Fig 2a) or a single-cell-derived HAP1 *SLC30A1* KO clone (Δ*SLC30A1*_1777_10) showed increased levels of zinc compared to control cell lines (Fig 2a, Suppl Fig 2b). This effect was reversed by ectopic expression of SLC30A1 (Fig 2a, Suppl Fig 2c-d) in the HAP1 SLC30A1-deficient cells. Moreover, a mutant SLC30A1, defective in its transport activity (D254A_H43A)^25^, failed to decrease the intracellular zinc levels (Fig 2a) despite localizing preferentially to the plasma membrane, similar to the WT protein (Suppl Fig 2c-d). These results therefore support the role of SLC30A1 as a zinc exporter. Zinc ions, together with Ca^2+^ and Mg^2+^, function as secondary messengers as well as cofactors in a variety of zinc metalloproteins, possibly affecting several cellular processes critical for the viral life cycle. Moreover, Zn^2+^ is directly implicated in regulating the activity of several pro-apoptotic caspases such as caspase-3, -6, -7, -8 and -9^26–28^. Since VSV is a cytolytic virus, able to kill infected cells via induction of apoptosis through both intrinsic and extrinsic pathways^12^, we speculated that elevated intracellular zinc levels might interfere with the activation of pro-apoptotic signalling through inhibition of caspases. To experimentally verify this hypothesis, we monitored the cleavage of different apoptotic caspases in VSV-infected A549 and HAP1 cells (Fig 2b-c). Indeed, A549 cells lacking the SLC30A1 transporter had impaired levels of all expressed and cleaved caspases tested (caspase-3, -7, -9) (Fig 2b). This effect was even more pronounced in the HAP1 *SLC30A1*-deficient clone (Fig 2c). Consistent with diminished caspase activation, we observed reduced number of Annexin V-positive *SLC30A1* KO cells compared to WT A549 cells (Fig 2d). In line with this data, we observed a reduction of Annexin V-positive cells upon VSV infection in WT cells supplemented with Zn^2+^ (ZnCl2 concentrations above 100mM) (Fig 2e). Altogether, these data show that loss of *SLC30A1* in A549 and HAP1 cells results in increased intracellular levels of Zn^2+^ and that cells lacking this transporter, upon VSV infection, show reduced caspases activation and apoptotic cell death.

**Fig. 2.**
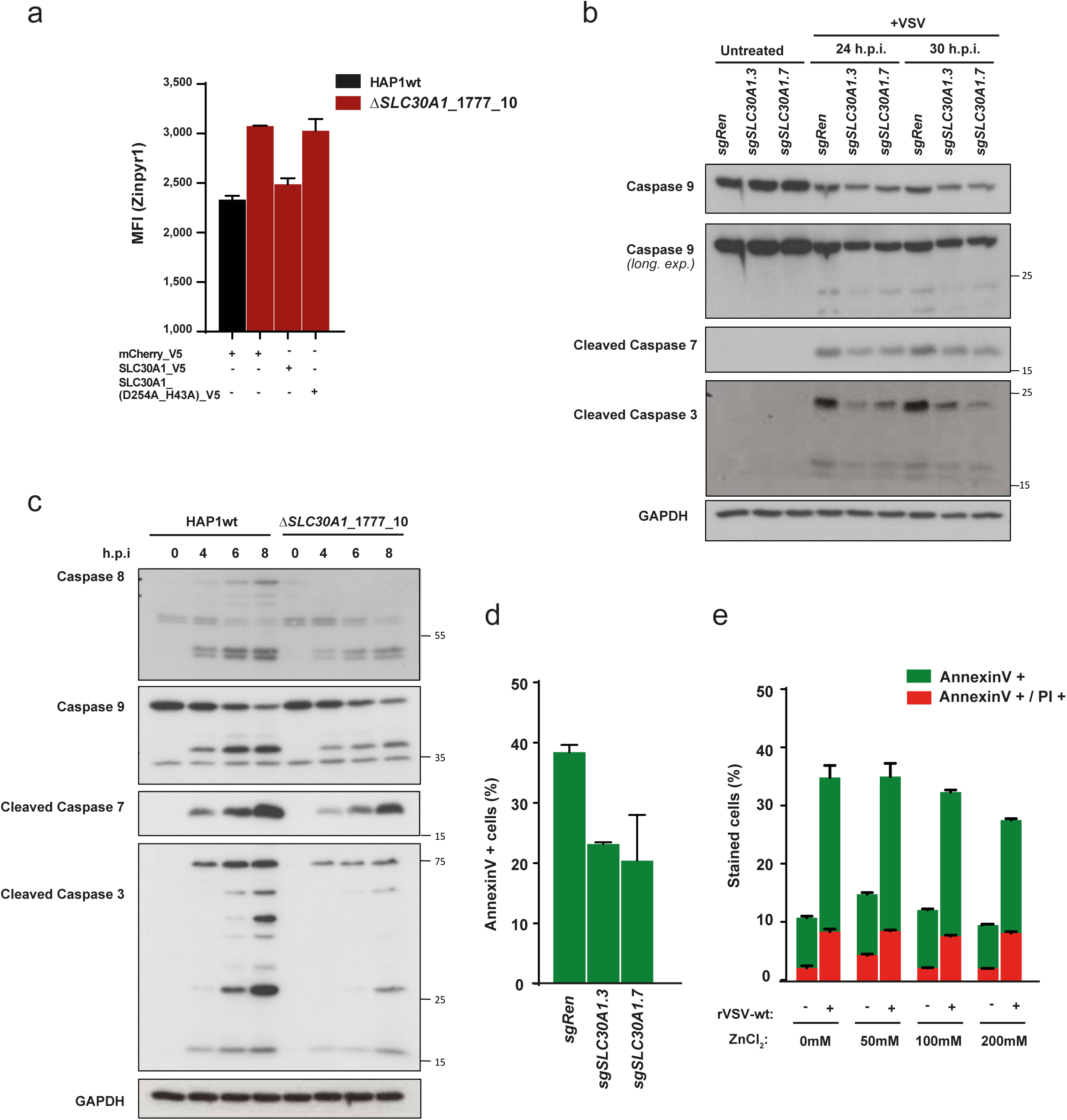
SLC30A1 inhibits apoptotic caspase activation via regulation of the intracellular Zn^2+^ levels. **a.** Intracellular Zn^2+^ levels were determined using Zinpyr1 fluorescent dye. **b-c.** Pro-apoptotic caspases activation of VSV infected A549 (**b.**) and HAP1 (**c.**) *SLC30A1* knock out cells. Cropped images are shown for conciseness. Full-length blots are presented in Supplementary Figure 5. **d**. Percentage of apoptotic cells in A549 cells expressing sgRNAs targeting *SLC30A1* or *Renilla* luciferase after infection with VSV (MOI 2) for 24 hours, as measured with AnnexinV-FITC. **e.** Quantification of AnnexinV and PI stained A549 WT cells pre-treated with indicated concentrations of ZnCl_2_ and infected with VSV-rWT (MOI 2) for 24 hours. Representative results from at least two independent experiments, flow cytometry experiments performed in triplicates.

### SLC30A1-deficient cells show reduced infectivity in multi cycle infections

We next tested whether loss of *SLC30A1* affected susceptibility to VSV infection in term of entry and or replication. Infection of HAP1 cells at high MOI did not result in differences in percentages of infected cells in *SLC30A1* HAP1 KO cells at early time points (Fig 3a), indicating that loss of this transporter did not impaired viral entry. However, upon low MOI VSV infection and at later time (which results in multi-cycle infections) we observed a strong reduction in the number of infected cells (Fig 3b). The reduction in infectivity associated with *SLC30A1* loss in HAP1 cells could be reverted by overexpression of the transport-competent form of SLC30A1. In contrast, the D254A_H43A transport mutant failed to rescue the viral phenotype, suggesting that the transport function was indeed crucially required (Fig 3b). The reduction in infectivity was further confirmed in independently generated populations of HAP1 and HEK293T, but not A549, cells transduced with different sgRNAs targeting *SLC30A1* (Fig 3c, Suppl Fig 3a-b). We did not observe any reduction in infectivity upon treatment with the pan-caspase inhibitor z-VAD-FMK, which reduces cleavage of caspase-3 to levels comparable to the *SLC30A1* knockout (Suppl Fig 3c-d), in line with previous observations suggesting that inhibition of caspases may temporarily attenuate VSV-induced apoptosis but does not actually affect virus replication^29,30^. Given the well described role of Zn^2+^ ions and other zinc transporters in the immune response against different pathogens and inflammatory stimuli^31^, we set to test whether *SLC30A1* loss may affect the antiviral immune response against VSV infection. Wild-type VSV virus does not generally elicit strong immune responses (Fig 3d). In contrast, VSV harbouring mutations in the matrix (M) protein create an attenuated form that fails to shut down host cell innate immune signalling efficiently and therefore elicits a stronger immune response. One of these mutants, VSV-ΔM51-GFP^32^, caused a readily measurable response upon infection of HAP1 WT cells at low MOI, as monitored by phosphorylation of IRF3 and STAT1 transcription factors (Fig 3d). Similar to WT VSV, loss of SLC30A1 resulted in reduced infectivity upon treatment with VSV-ΔM51-GFP (Suppl Fig 3e). However, cells deficient for *SLC30A1* did not show notable differences in the activation of IFNα/β and NF-κB pathways, two of the major antiviral immune response pathways, compared to wild type cells (Fig 3d). Moreover, we did not detect differences at early hours post infection in either accumulation of viral RNA (0.5-4h) or viral protein (i.e. VSV-G, 6h) (Fig 3e-f). Interestingly, when measuring the titers of virus produced by the *SLC30A1* knockout cells compared to HAP1 WT we did measure a strong reduction in the number of viral particles released from the *SLC30A1*-deficient cells (Fig 3g). This is consistent with the reduced amount of viral RNA in the *SLC30A1*-deficient cells at the later stages of the infection (≥4 hours), when the reinfection with the produced viral particles starts to occur (Fig 3e). Together, these observations suggest that SLC30A1-deficiency affects VSV at late stages of the virus life cycle. Overall, we did not observe effects on cell infectivity or immune response due to *SLC30A1*-loss under single-cycle conditions (high MOI). Interestingly, *SLC30A1*-deficient cells produced reduced titers of virus after infection, likely to be due to a defect at the late stage of the virus cycle, which could become relevant in a multicycle experiments.

**Fig. 3.**
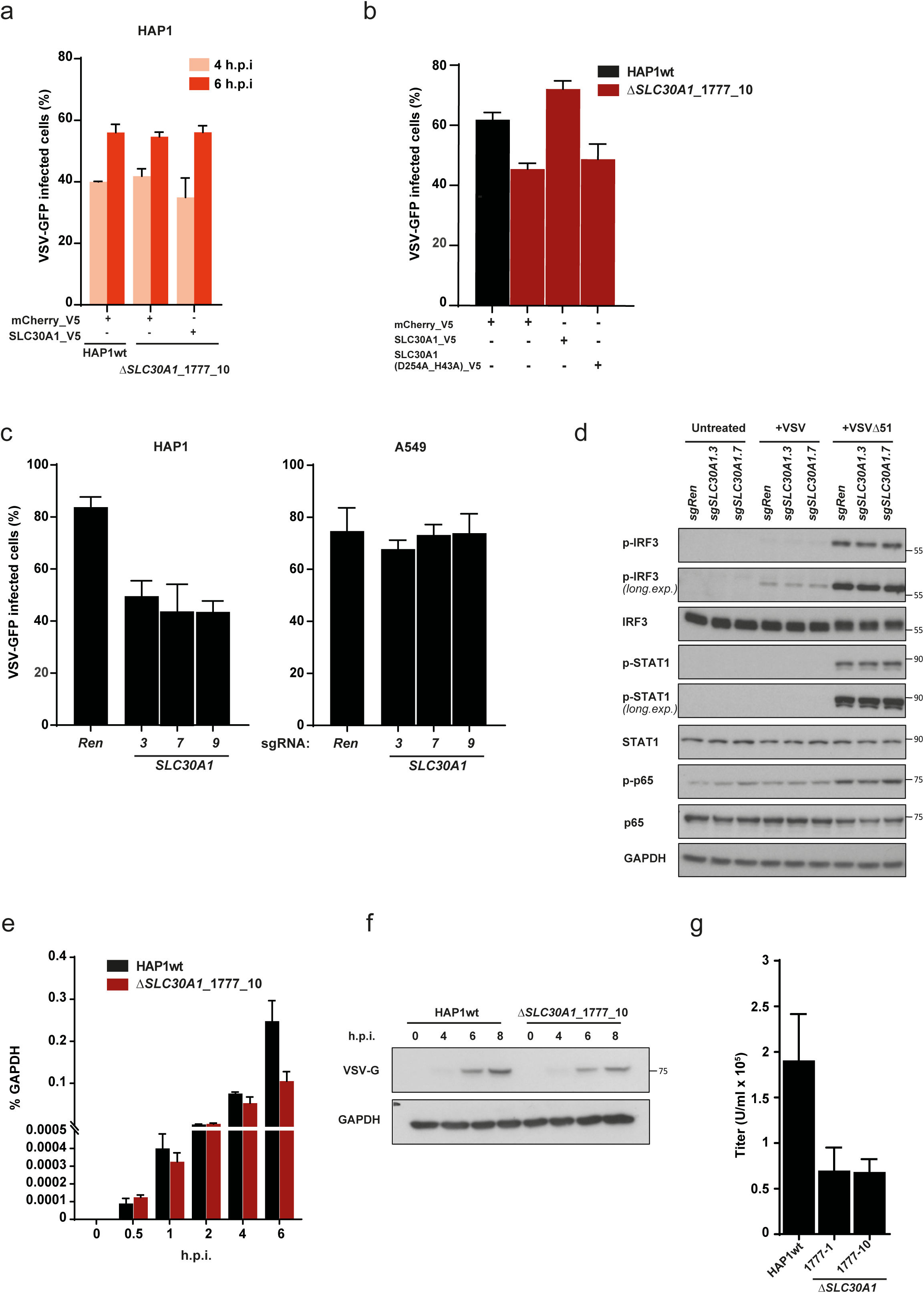
SLC30A1 loss affects VSV infection through modulation of Zn levels. **a.** to **c.** VSV-GFP virus replication in *SLC30A1* knock-out A549 and HAP1 cells and HAP1 cells overexpressing SLC30A1wt and SLC30A1(D254A_H43A) cDNAs infected with MOI 1 (**a**) and MOI 0.001 and 0.002, (**b** and **c**) respectively. The number of GFP+ cells was quantified with flow cytometry at 14 h.p.i. (**b** and **c**) and indicated time points (**a**). **d.** Immunoblot analysis of type I IFN/STAT1 signalling pathway activation in the VSV-infected (MOI 2) HAP1 cells carrying sgRNAs against *SLC30A1* and *Renilla* luciferase control. Cropped images are shown for conciseness. Full-length blots are presented in Supplementary Figure 5. **e.** Quantitative RT-PCR of total viral RNA in the HAP1 wild type cells and a *SLC30A1* single cell KO clone at different points after VSV infection (MOI 2). **f.** VSV-G protein levels in the HAP1 wild type cells and SLC30A1 single cell KO clone at different points after VSV infection (MOI 2). Cropped images are shown for conciseness. Full-length blots are presented in Supplementary Figure 5. **g.** Quantification (flow cytometry) of virus titer produced by HAP1wt and two *SLC30A1* single cell KO clones infected at MOI of 2 with VSV-GFP for 8 hours.

### Loss of SLC35A1 results in increased cell death and apoptosis upon VSV infection

The sialic acid-CMP transporter SLC35A1 has been previously shown to be involved in the cellular response to apoptotic stimuli. In particular, *SLC35A1* loss has been reported to evoke an elevated Golgi stress response^31^ resulting in increased rates of pro-apoptotic signalling in the cells^32–34^. We therefore hypothesised that loss of SLC35A1 may result in sensitisation to cell death induction upon VSV infection as well as upon other stimuli. Indeed, we observed that A549 and HAP1 cells lacking SLC35A1 (Suppl Fig 4a) show an elevated cell death and apoptotic response in response to VSV infection, as monitored by increased number of Annexin V-positive cells (Fig 1f, 4a, Suppl Fig 1j) and increased caspase cleavage that could be partially reversed by ectopic overexpression of *SLC35A1* (Suppl Fig 4b-c) suggesting a dependency on SLC function. Cell death was similarly observed upon stimulation with compounds previously reported to induce a Golgi stress response such as Brefeldin A^35^ (Suppl Fig 4d). However, in cells lacking *SLC35A1* we did not observe an increase in MAPK pathway activity or ARF4 protein previously reported to indicate a response to Golgi stress (Fig 4b)^35^, suggesting involvement of alternative pathway(s). Consistent with this, we found that *SLC35A1* knockout cells were also more sensitive to two additional, non-Golgi stress-related, cytotoxic compounds: the topoisomerase inhibitor camptothecin and the proteasome inhibitor carfilzomib (Suppl Fig 4d). Interestingly, *SLC35A1* KO cells show increased infection rates in multi-cycle infections in several different cell lines (Fig 4c, Suppl Fig 4e). This effect could also be reverted by re-expression of *SLC35A1* (Fig 4d). Finally, in experiments following a single infection cycle, we detected increased virus titers in *SLC35A1* KO cells compared to wild type cells as measured by FACS (Fig 4e). Altogether, these data point to a combination of a more pronounced apoptotic response and an increased rate of VSV replication, as indicated by the higher virus titers produced in cells lacking *SLC35A1*, as factors contributing to its role in cellular survival upon viral infection.

**Fig. 4.**
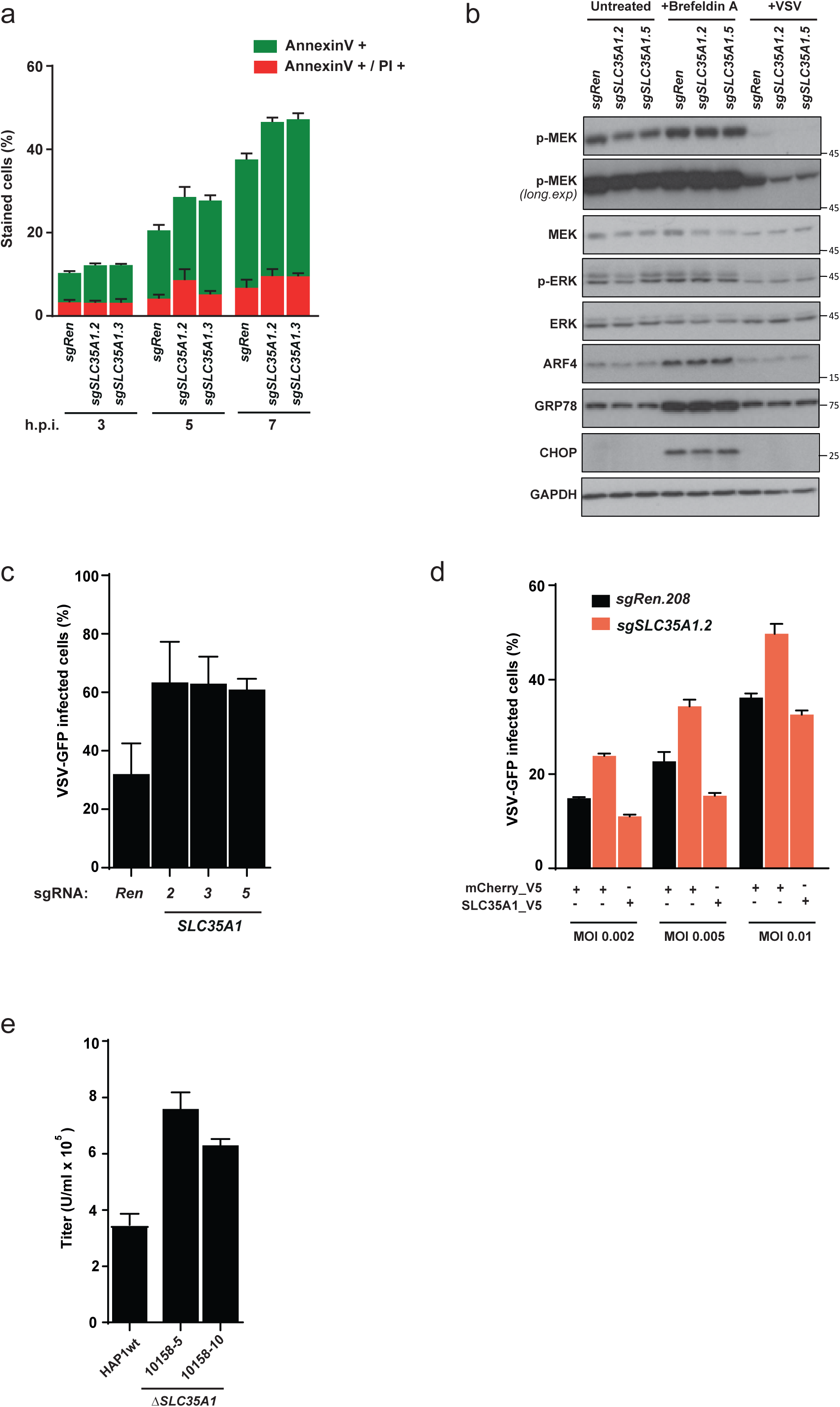
SLC35A1 loss promotes cell death and apoptosis upon VSV infection. **a.** Percentage of apoptotic (AnnexinV+) and necrotic (AnnexinV+/PI+) cells in the HAP1 cell line expressing sgRNAs against *SLC35A1* and *Renilla* luciferase after infection with VSV at MOI 2. **b**. Immunoblot analysis of Golgi stress signalling pathway activation in the VSV-infected (MOI 2) and Brefeldin A-stimulated (3?g/ml) A549 cells carrying sgRNAs against *SLC35A1* and *Renilla* luciferase. Cropped images are shown for conciseness. Full-length blots are presented in Supplementary Figure 5. **c-d.** VSV-GFP virus replication in *SLC35A1* knock out HAP1 cells (**c**) and HAP1 cells overexpressing SLC35A1 cDNA (**d**) infected with MOI ranging from 0,001 to 0,01 for 14 hours. **e.** Quantification (flow cytometry) of virus titer produced by HAP1wt and two *SLC35A*1 single cell knock out clones infected with MOI of 2 VSV-GFP for 8 hours. Representative results from at least two independent experiments, flow cytometry experiments performed in triplicates.

## Discussion

Functional genomic and genetic screens, based on siRNA, insertional mutagenesis and, more recently, CRISPR/Cas9, have been extremely valuable in identifying host factors affecting viral infectivity and replication, in particular regarding entry factors^16,17,36–38^. However, the role of metabolic genes and transporters as host genes affecting the viral life cycle and host cell survival upon infection remain relatively understudied. Here we employed SLC-focused CRISPR knockout screens to identify two solute carriers, the CMP-sialic acid/CMP transporter *SLC35A1*/CST and the zinc transporter *SLC30A1*/ZnT1 as critical host factors affecting in opposite ways the survival of human cell lines towards infection with the oncolytic VSV virus (Fig 1). We confirmed that SLC30A1 is a zinc exporter localized at the plasma membrane and that genetic inactivation resulted in increased intracellular zinc levels and increased resistance to VSV-induced cellular killing (Fig 1-2). Mechanistically, loss of *SLC30A1* did not appear to affect the anti-viral immune response or viral entry (Fig3). We instead observed a reduced infection of *SLC30A1*-deficient cells compared to WT cells in multi-cycle infections. This phenotype was coupled to reduced virus titers, potentially hinting towards a defect in the later stages of the viral cycle (Fig3). More importantly, upon *SLC30A1* inactivation we observed a marked reduction of caspase activation and, consequently, reduced apoptotic cell death (Fig 2). Given the known role of Zn^2+^ ions as pan-caspase inhibitors^26–28^, we hypothesise that loss of SLC30A1, and the resulting increased intracellular Zn^2+^ levels, result in an apoptosis-resistant state and an accompanying increased survival upon further perturbation such as VSV infection.

The second transporter gene identified in our VSV screen, the Golgi-residing SLC35A1, has previously been implicated in IAV infectivity due to its critical role in the addition of sialic acid, the major receptor of influenza virus, to proteins and lipids on the cell surface^39,40^. When we screened for solute carriers affecting infectivity to IAV, we identified not only *SLC35A1* but also the related transporter *SLC35A2*, a UDP-galactose/UDP transporter, as hits (Fig 1). Both genes were separately identified in genome-wide screens^16,17^ as host factors required for IAV infectivity and recapitulated in our screen. Unlike the observation with IAV, loss of *SLC35A1* resulted in increased apoptotic cell death upon VSV infection (Fig 1, 4). However, we did not observe activation of signalling pathways commonly associated with the induction of Golgi stress, a pro-apoptotic pathway previously shown to be linked to SLC35A1^31^. It was formerly described that sialic acid modifications play important roles for various pro-apoptotic signals^41–43^ as well as prevent apoptosis induced by galectins^42^. Further studies will be required to characterize which pathway(s) are involved in VSV- and SLC35A1-dependent cell death and viral phenotypes.

In conclusion, we used an unbiased SLC-focused genetic approach to identify transporter proteins affecting the survival of human cancer cell lines upon VSV infection. Both genes identified, *SLC30A1* and *SLC35A1*, represent novel host factors that affected the induction of apoptosis upon viral infection, albeit with opposite roles. Taken together, these results offer new insights and may spark new ideas into possible strategies to pharmacologically interfere with viral infections, especially in the context of oncolytic viruses.

## Methods

### Cell lines and viruses

HEK293T and A549 cells were obtained from ATCC (Manassas, VA, USA), MDCK2 cells were kindly provided by G.Versteeg (MFPL, University of Vienna, Austria). HAP1 were obtained from Horizon Genomics. Cells were cultured in DMEM or IMDM medium supplemented with 10% (v/v) FBS and antibiotics (100 U/mL penicillin and 100 mg/mL streptomycin) (Gibco). Cell lines were routinely checked for mycoplasma using MycoAlert kit (Lonza). The virus strains used in this study were recombinant VSV-GFP^44^, VSV-rWT^45^, VSV-ΔM51-GFP^46^ and IAV-WSN/1933 (H1N1). Viruses were grown on MDCK2 and Vero cells (both grown in DMEM supplemented with 10% FBS and antibiotics) and titrated using plaque assay. SLC35A1 and SLC30A1 single cell knockout HAP1 clones were obtained from Haplogen Genomics (SLC30A1 clones: 1777-1 and 1777-10, SLC35A1 KO clones: 10158-5, 10158-10).

### CRISPR-KO screening

The human SLC knockout CRISPR/Cas9 library been described previously^15^. Viral particles were prepared by transient transfection of low passage, subconfluent HEK293T cells with the library plasmid pool together with packaging plasmids psPAX2 (Addgene #12260) and pMD2.G (Addgene #12259) using PolyFect (Qiagen). After 24 hours the media was changed to fresh RPMI media supplemented with 10% FCS and antibiotics. The viral supernatant was collected after 48 hours, filtered (0.45 μm) and stored at −80°C until further use.

A549 cells were infected with the SLC knockout library at a multiplicity of infection of 0.2-0.3 and screened after selection for 5 days with puromycin (2 μg/mL) and 2 days of recovery and expansion. For the screen, 8×10^6^ (equals ∼2000x coverage) of A549 cells were seeded in duplicates and infected with IAV-WSN and VSV-GFP at the MOI of 0.5 and 10 respectively. Cells were collected 96 hours post infection (for the IAV-WSN screen) and at 48h hours post infection (for the VSV-GFP screen) together with time matching uninfected controls. Additionally, time 0 samples were collected at the time of infection. Genomic DNA was extracted using the DNAeasy kit (Qiagen) and amplified with 2 rounds of PCR using primers derived from Shalem et al.^47^ and modified to allow dual indexing by addition of an additional barcode on the forward primers. The PCR-amplified samples were sequenced on a HiSeq2500 (Illumina) at the Biomedical Sequencing Facility (CeMM/MUW).

### Screen analysis

Sequencing reads were matched to the sgRNA library sequences and counted using an in-house script. One sample was removed from the analysis because of low coverage (median of less than 50 reads per sgRNA). Guide-level read tables were input to a gene-level DESeq2 model^48^. Custom normalization factors were provided assuring equal median sgRNA read per sample. As the gene-level model included the effect of the guides, identical number of guides (i.e. 6) per gene were required across all genes. Therefore, for 10 genes with only 5 guides in the library, a randomly selected guide was duplicated, and six guides were randomly selected for 3 genes with more than 6 guides in the library as well as the for the negative control. The model was tested for differential abundance contrasting virus-infected samples versus the uninfected control samples of corresponding time points. Genes with a multiple-testing adjusted p-value^49^ of less than 0.1 (adj. p-value <0.1) were considered significantly enriched or depleted, corresponding to an estimated false-discovery rate below 10%.

### Plasmids and knockout cell lines generation

For CRISPR-based knockout cell lines, sgRNAs were designed using CHOPCHOP^50^ and cloned into pLentiCRISPRv2 (Addgene, #52961), pLentiCRISPRv3^51^, pX459, LGPIG (pLentiGuide-PuroR-IRES-GFP) or LGPIC (pLentiGuide-PuroR-IRES-mCherry)^52^. sg*Ren* targeting *Renilla luciferase* cDNA was used as negative control sgRNA^52^(Suppl Table 2). SLC35A1 (HsCD00415788) and SLC30A1 (HsCD00375357) cDNAs were obtained as gateway-compatible pENTR vectors from the Harvard PlasmID Repository. sgRNA-resistant cDNA versions as well as the SLC30A1 D254A_H43A double mutant^25^ were generated using NEB Q5 site-directed mutagenesis kit (Suppl Table 2). cDNAs were transferred into gateway-compatible lentiviral expression vectors pLX304 (Addgene, #25890, CMV promoter driven expresion) or pRRL (described previously^52^, EF1a promotor driven expression) using LR recombination (ThermoFisher Scientific).

For generation of lentiviral knockout and overexpression cells HEK293T cells were transfected with psPAX2 (Addgene #12260) and pMD2.G (Addgene #12259) and expression vectors using Polyfect (Qiagen). 24 hours post transfection medium was replaced with fresh medium that was harvested 48 hours later, filtered (0.45 μm), supplemented with 5μg/mL Polybrene (Hexadimethrine bromide, Sigma) and added to target cells. 48 hours after transduction the medium was supplemented with the respective selection antibiotics. Cells were selected for 5-7 days. Due to the downregulation of the expression of wild type cDNA of SLC30A1 over time, fresh overexpression cells were prepared every 2-3 weeks. Editing efficiency of sgRNAs was assessed by Sanger sequencing followed by analysis of sequencing results with TIDE web tool^53^.

### Flow cytometry

For flow cytometric analyses of virus-infected cells, cells were seeded into 24-well plates and infected with IAV or VSV at the specified MOIs. Cells were collected at indicated time points and fixed with 4% PFA in PBS. IAV-WSN/1933 infected cells were permeabilized (PBS, 0.1% Triton X-100), stained with AlexaFluor-488-labelled (ThermoFisher Scientific) anti-influenza nuclear protein antibody (ab20343, Abcam) for 1 hour at room temperature followed by two washing steps.

In order to determine the amount of dead or apoptotic cells after virus infection, cells were seeded into 12-well plates, infected at specified MOIs, and at the indicated time points. Depending on the cells prepared for staining, cells were stained with AnnexinV-FITC (eBioscience) or AnnexinV-AF647 (eBioscience), Live/Dead Red or Green fixable dyes (Invitrogen) or Propidium Iodide (PI) for 15min (AnnexinV and PI) or 30min (Live/Dead) at room temperature, fixed with 4% PFA and subsequently analysed by flow cytometry.

For the measurement of intracellular Zn^2+^ levels cells were incubated for 30min at 37°C in serum-free IMDM supplemented with 2mM EDTA and 50μM Zipyr 1 (Abcam, ab145349).

Flow cytometry-based multi-color competition assay (MCA) was performed as described previously^52^. Briefly, A549 cells expressing LGPIC-sgRen were mixed in 1:1 ration with LGPIG reporter cells containing sgRNAs targeting the gene of interest. Mixed cell populations were infected with VSV-rWT at MOI 5. The respective percentage of viable (FSC/SSC) mCherry-positive and eGFP-positive cells at the indicated time points was quantified by flow cytometry. Samples were analysed on an LSR Fortessa (BD Biosciences) and data analysis was performed using FlowJo software (Tree Star Inc., USA).

### RNA isolation and qRT-PCR

For qRT-PCR measurement of viral RNA (vRNA), cells were infected with VSV-GFP at MOI 20 and at the indicated time points total RNA was isolated from the samples using RNeasy Kit (Qiagen). RNA was reverse transcribed using random hexamer primers and RevertAid Reverse Transcriptase (Fermentas). qRT-PCR was performed using SensiMix SYBR Green (Bioline) on a QIAGEN Rotor-Gene Q. Results were normalized to the levels of the housekeeping gene *GAPDH* (Suppl. Table 2).

### Viral titer measurement

For the measurement of virus progeny, cells were infected with VSV-GFP at MOI of 2. After incubation with infectious media for 30min at 37°C, cells were subsequently washed with PBS and media replaced with full IMDM for the remaining duration of the experiment (8 hours in total). Media, containing viral particles, was serially diluted and added to fresh HAP1 wild type cells. Cells were further incubated for 5 hours and the percentage of infected cells was analysed by flow cytometry. Viral titer was calculated using the following formula: U/mL = (cell number * % of GFP^+^ cells) / (volume of virus containing media * dilution factor).

### Antibodies and Immunoblotting

The following antibodies were used: V5 (Invitrogen, R960-25,) GAPDH (Santa Cruz, sc-365062), alpha tubulin (Abcam, ab-7291-100), pIRF3 (Cell signalling, #4947), IRF3 (Cell signalling, #11904), pSTAT1 (Cell signalling, #9171), STAT1 (BD Transduction, #610115), p-p65 (Cell signalling #3033), p65 (Cell signalling #8242), cleaved caspase 3 (Cell signalling #9661), cleaved caspase 7 (Cell signalling #9491), caspase 8 (Cell signalling #9746), caspase 9 (Cell signalling #9502), phospho-ERK1/2 T202/Y204 (Cell signalling #4370), ERK1/2, (#4694, Cell Signaling), phospho-MEK1/2 S217/221 (#9154, Cell Signaling), MEK1/2 (#9126, Cell Signaling), ARF4 (#11673-1-AP, Proteintech), CHOP (#MA1-250, Invitrogen), GRP78/BiP (#610979, BD Biosciences) and VSV-G (8G5F11, Kerafast). The following secondary antibodies were used: goat anti-mouse HRP (115-035-003, Jackson ImmunoResearch) and goat anti-rabbit HRP (111-035-003, Jackson ImmunoResearch).

For immunoblotting whole cell extracts were prepared using RIPA lysis buffer (25mM Tris/HCl pH 7.6, 150mM NaCl, 1% NP-40, 1% sodium deoxycholate, 0.1% SDS and one tablet of EDTA-free protease inhibitor (Roche) per 50 mL) supplemented with Halt phosphatase inhibitor cocktail (Thermo Fisher Scientific #78420) and Benzonase (71205, EMD Millipore). Protein extracts were normalized using the Bradford assay (Bio-Rad). Cell lysates were run on SDS-polyacrylamide gel and transferred to nitrocellulose membranes Protran BA 85 (GE Healthcare). The membranes were incubated with the antibodies indicated above and visualized with horseradish peroxidase–conjugated secondary antibodies using the ECL Western blotting system (Thermo Scientific).

### Immunofluorescence

For immunofluorescence detection of subcellular localisation of overexpressed SLC30A1 WT and mutant fused to mScarlet in HAP1 wild type cells, cells were seeded onto poly-L-lysine hydrobromide (P6282, Sigma-Aldrich)-coated 96-well CellCarrier Ultra plates (PerkinElmer). After an attachment period, cells were stained for the measurement of the intracellular Zn^2+^ levels as described above. Images were acquired on the Opera Phenix automated spinning disk confocal microscope (PerkinElmer).

## Acknowledgments

We would like to thank Gijs Versteeg (University of Vienna), Michael Freissmuth (Medical University of Vienna) and all the members of the Superti-Furga laboratory for critical input. In particular, we would like to thank Johannes W. Bigenzahn and Manuele Rebsamen for valuable suggestions and scientific insights as well as Bojan Villagos for graphical design. CeMM and the Superti-Furga laboratory are supported by the Austrian Academy of Sciences. We acknowledge receipt of third-party funds from the Austrian Science Fund (FWF W1205 DK CCHD, A.M., FWF P29250-B30 VITRA, E.G, J.K, G.F., FWF F4711-B20 Myeloid Neoplasms. F. K.), the European Research Council (ERC AdG 695214 GameofGates, A.M., U.G.), the European Commission (Marie Sk1odowska-Curie Action Fellowship 661491, E.G).

## Author contributions

A.M., E.G and G.S-F. conceived the study. A.M. performed experiments and analysed the data. U.G. analysed data. F.K., S.L., J.K and G.F. performed experiments and/or provided reagents or technical know-how. A.M., E.G. and G.S-F. wrote the manuscript.

## Additional information

The authors declare no competing interests.

## Data availability

The datasets generated during and/or analysed during the current study are available from the corresponding author on reasonable request.

## Supplementary Figures

**Suppl.Fig. 1 Genetic screens for cell survival upon IAV and VSV infection. a.** Violin plots of raw and normalized read counts and **b.** Correlation of normalized read counts across biological replicates in the IAV-WSN screen. Quantification of flow cytometry-based viral replication assay for HAP1 *SLC35A1* **(c),** *SLC35A2* **(d)** knockout cell lines and cells re-expressing SLC35A1 cDNA **(e)**. Cells were infected with IAV-WSN/33 at MOI 0,01. At 24 h.p.i. the number of the infected cells was assessed after staining with anti-Influenza Nuclear Protein antibody coupled to AF488. Representative results from at least two independent experiments performed in triplicates. **f.** Assessment of A549 cell death upon infection with wild type VSV-GFP (MOI of 10). Cells were incubated for the indicated time points and relative cell number was determined using crystal violet staining. **g.** Violin plots of raw and normalized read counts and **h.** Correlation of normalized read counts across biological replicates in the VSV-GFP screen. A549 cells expressing sgRNAs targeting SLC30A1 (**i**) or *SLC35A1* (**j**) were infected with wt VSV-GFP at MOI of 5 and at the indicated time points stained with Live/Dead™ cell viability dye. Positive cells were quantified with flow cytometry.

**Suppl.Fig. 2. SLC30A1 loss results in increased intracellular Zn^2+^ concentration and decreased virus-induced cell death. a.** Editing efficiency of sgRNAs targeting the *SLC30A1* gene as quantified by the TIDE sequencing method in A549 cells. **b**. Representative FACS histogram and quantification of the A549 SLC30A1 knock out cells stained with Zn^2+^ reactive dye – Zinpyr1. **c.** Representative immunoblot of V5 tagged wtSLC30A1 and SLC30A1(D254A_H43A) double mutant cDNA expressing cells. Cropped images are shown for conciseness. Full-length blots are presented in Supplementary Figure 5. **d.** Immunofluorescent image of HAP1wt cells overexpressing mScarlet-fused wild type SLC30A1 and SLC30A1(D254A_H43A) double mutant stained with Zn^2+^ reactive fluorescent dye Zinpyr1.

**Suppl.Fig. 3. SLC30A1 in apoptosis and VSV infectivity. a.** Editing efficiency of sgRNAs targeting the *SLC30A1* gene as quantified by the TIDE sequencing method in HAP1 cells. **b.** Flow cytometry-based VSV-GFP virus replication in *SLC30A1* knock out HEK293T cells (MOI 0.001). **c.** VSV-GFP virus replication in *SLC30A1* KO HAP1 cells treated with caspase inhibitor z-VAD infected at MOI of Number of GFP+ cells was quantified with flow cytometry at 14 h.p.i. **d.** Immunoblot analysis of caspase 3 and 7 cleavage in HAP1wt cells stimulated with zVAD and SLC30A1 knock out clone upon infection with VSV-GFP (MOI 2). Cropped images are shown for conciseness. Full-length blots are presented in Supplementary Figure 5. **d.** VSV-Δ51-GFP virus replication in the HAP1 SLC30A1 knock-out and wild type cells infected with MOI 0.001 and 0.005 for 14 hours.

**Suppl.Fig. 4. SLC35A1 in apoptosis and VSV infectivity. a.** Editing efficiency of sgRNAs targeting *SLC35A1* gene as quantified using the TIDE sequencing method in HAP1 and A549 cells. **b.** Pro-apoptotic caspases activation of VSV infected HAP1 *SLC35A1* knock out cells and SLC35A1 cDNA-expressing cells. Cropped images are shown for conciseness. Full-length blots are presented in Supplementary Figure 5. **c.** Representative immunoblot of HAP1 cells expressing V5-tagged wild type SLC35A1. Cropped images are shown for conciseness. Full-length blots are presented in Supplementary Figure 5. **d.** Percentage of apoptotic (AnnexinV+) and necrotic (AnnexinV+/PI+) cells in the HAP1 cell line expressing sgRNAs against SLC35A1 and *Renilla* luciferase and stimulation with Brefeldin A (12μg/ml), Carfilzomib (4μM) and Campthotesin (2μM) for 8 hours. **e.** Flow cytometry based VSV-GFP virus replication in *SLC35A1* KO HEK293T cells (MOI 0.001).

**Suppl.Fig. 5. Full blots used in this study.**

## Tables

- **Suppl Table1**: Guide RNA read count tables from SLC KO screens
- **Suppl Table2**: Primers (qPCR, sequencing) and sgRNAs used in the study

