## Supplementary Figures for "The transporters SLC35A1 and SLC30A1 play opposite roles in cell survival upon VSV virus infection"

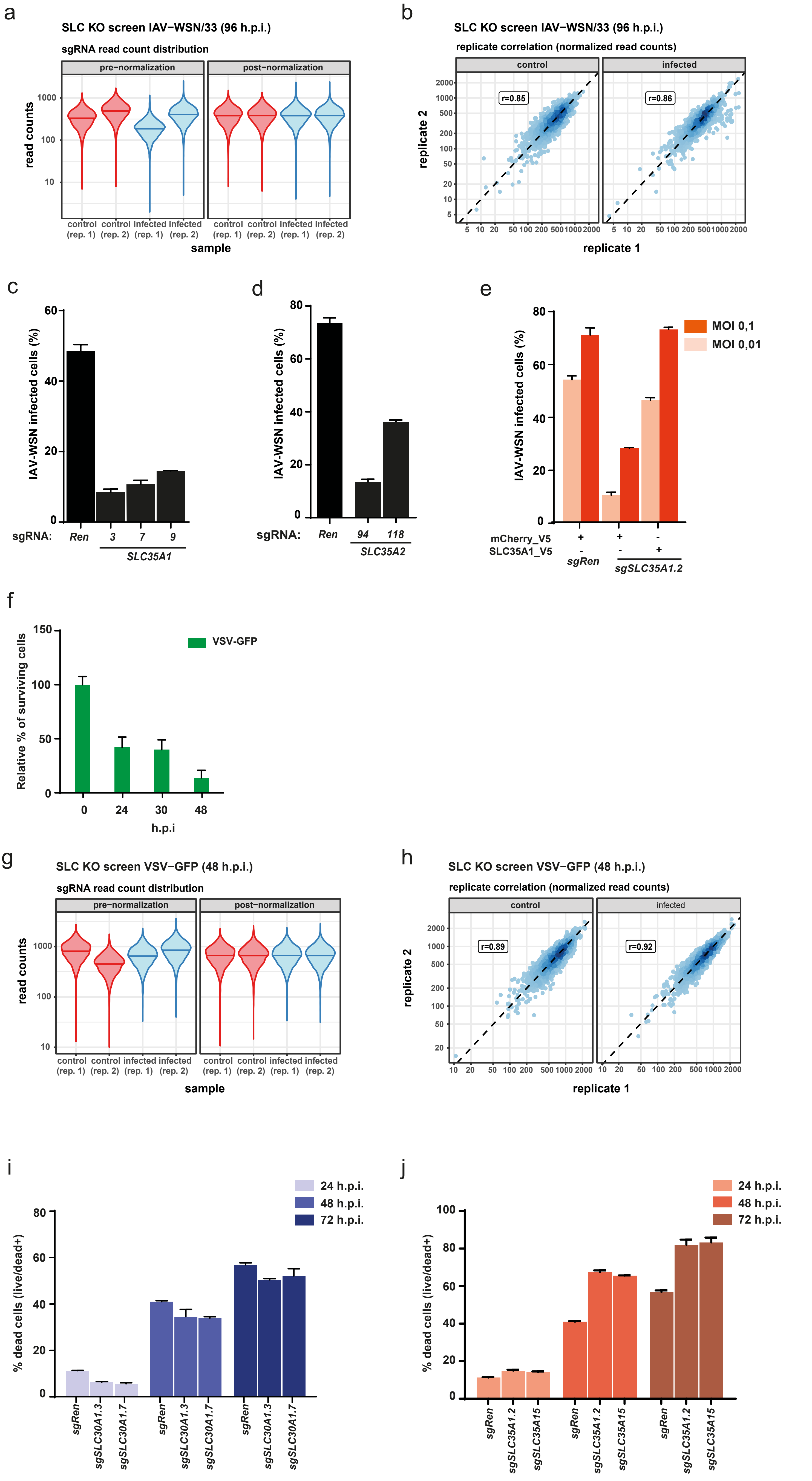
a
SLC KO screen IAV−WSN/33 (96 h.p.i.)
sgRNA read count distribution
pre−normalization
post−normalization
read counts
control (rep. 1) control (rep. 2) infected (rep. 1) infected (rep. 2)
sample
b
SLC KO screen IAV−WSN/33 (96 h.p.i.)
replicate correlation (normalized read counts)
control
infected
r=0.85
r=0.86
replicate 2
replicate 1
c
IAV-WSN infected cells (%)
sgRNA: Ren 3 7 9
SLC35A1
d
IAV-WSN infected cells (%)
sgRNA: Ren 94 118
SLC35A2
e
MOI 0,1
MOI 0,01
IAV-WSN infected cells (%)
mCherry_V5 + + -
SLC35A1_V5 - - +
sgRen
sgSLC35A1.2
f
VSV-GFP
Relative % of surviving cells
0 24 30 48
h.p.i
g
SLC KO screen VSV−GFP (48 h.p.i.)
sgRNA read count distribution
pre−normalization
post−normalization
read counts
control (rep. 1) control (rep. 2) infected (rep. 1) infected (rep. 2)
sample
h
SLC KO screen VSV−GFP (48 h.p.i.)
replicate correlation (normalized read counts)
control
infected
r=0.89
r=0.92
replicate 2
replicate 1
i
24 h.p.i.
48 h.p.i.
72 h.p.i.
% dead cells (live/dead+)
sgRen sgSLC30A1.3 sgSLC30A1.7
j
24 h.p.i.
48 h.p.i.
72 h.p.i.
% dead cells (live/dead+)
sgRen sgSLC35A1.2 sgSLC35A15

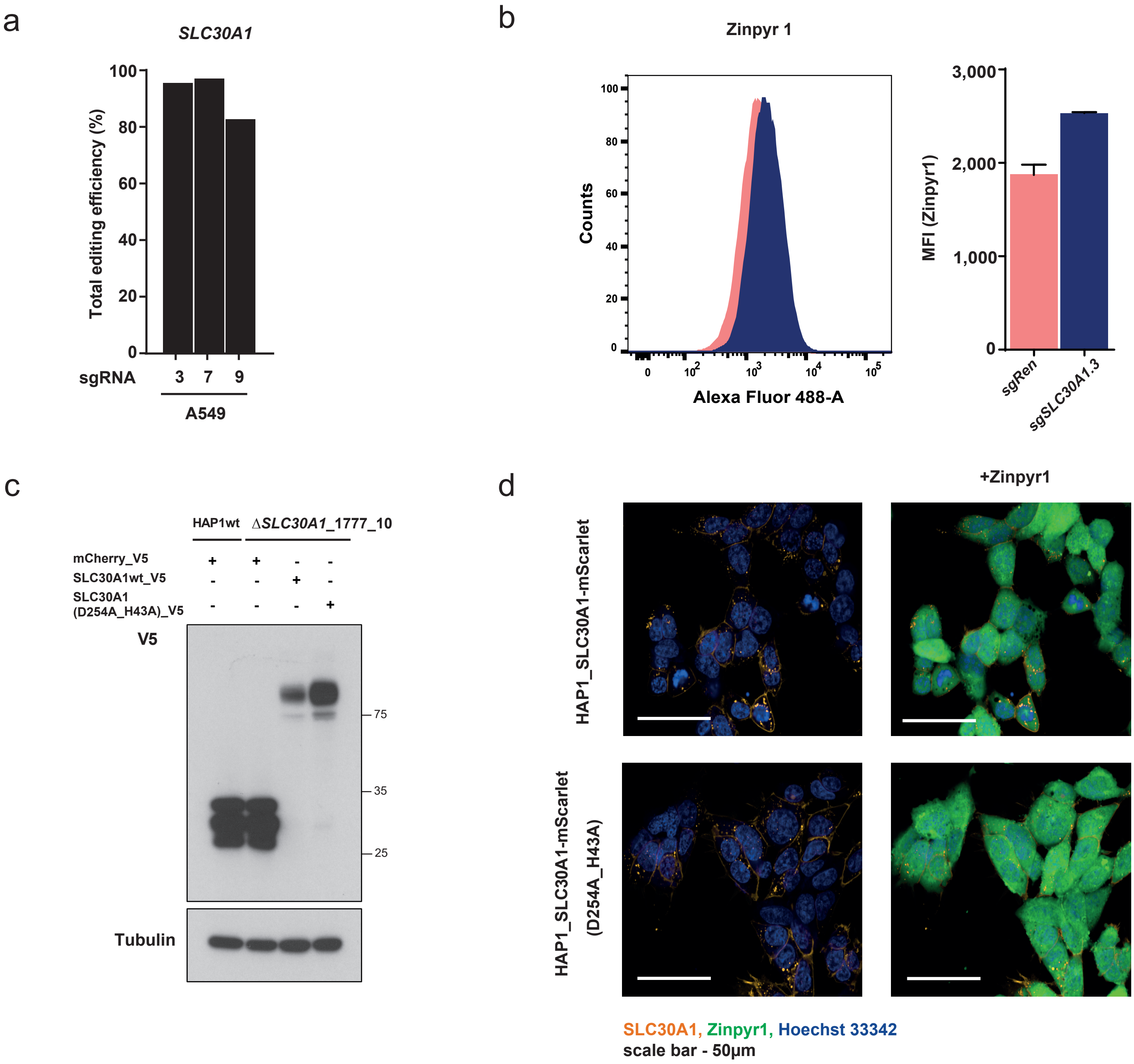
a
SLC30A1
Total editing efficiency (%)
100
80
60
40
20
0
sgRNA
3
7
9
A549
b
Zinpyr 1
Counts
Alexa Fluor 488-A
MFI (Zinpyr1)
3,000
2,000
1,000
0
sgRen
sgSLC30A1.3
c
HAP1wt
ΔSLC30A1_1777_10
mCherry_V5
SLC30A1wt_V5
SLC30A1
(D254A_H43A)_V5
V5
75
35
25
Tubulin
d
+Zinpyr1
HAP1_SLC30A1-mScarlet
HAP1_SLC30A1-mScarlet
(D254A_H43A)
SLC30A1, Zinpyr1, Hoechst 33342
scale bar - 50µm

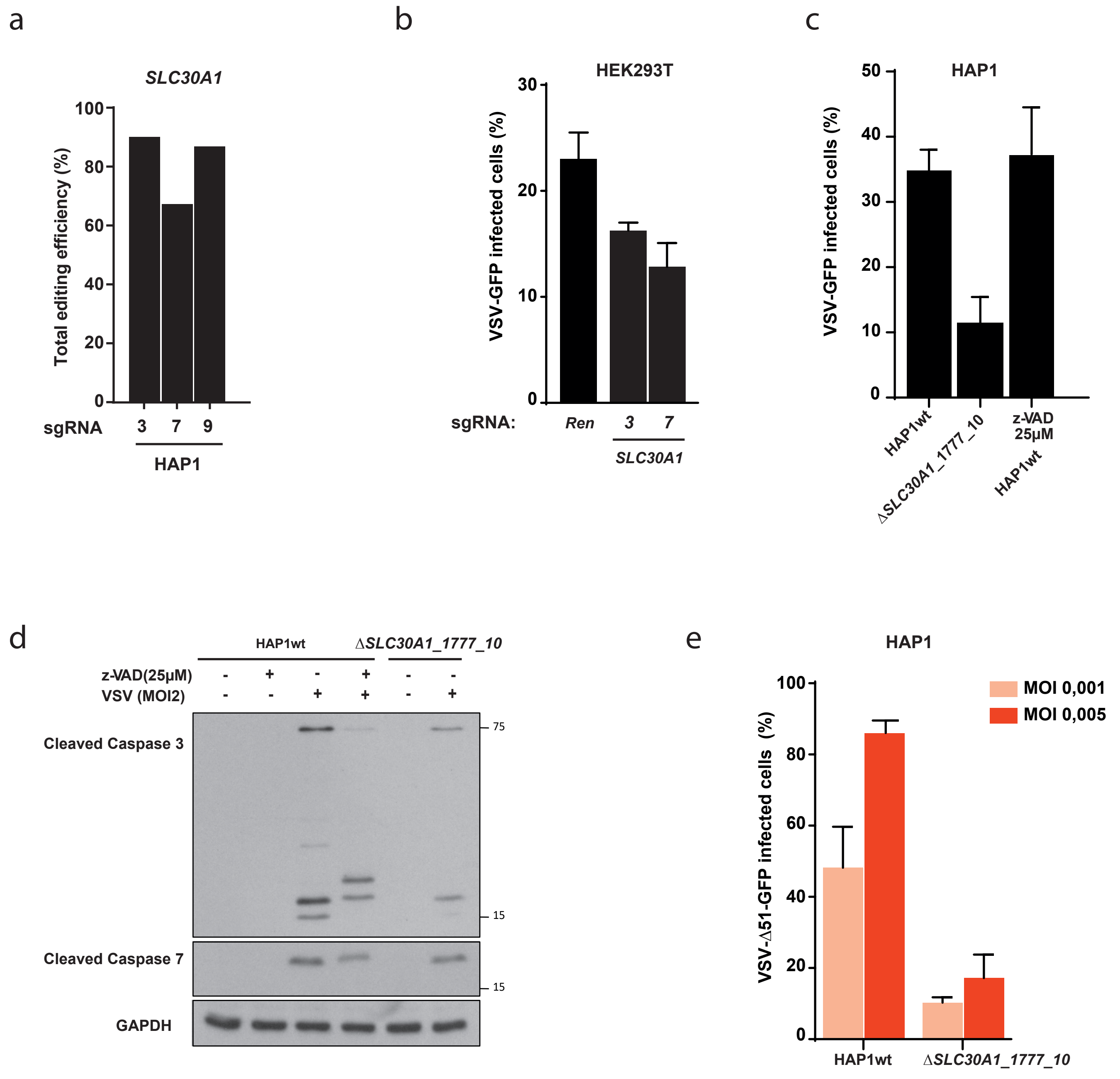
a
SLC30A1
Total editing efficiency (%)
100
80
60
40
20
0
sgRNA
3
7
9
HAP1
b
HEK293T
VSV-GFP infected cells (%)
30
20
10
0
sgRNA:
Ren
3
7
SLC30A1
c
HAP1
VSV-GFP infected cells (%)
50
40
30
20
10
0
HAP1wt
ΔSLC30A1_1777_10
z-VAD 25µM
HAP1wt
d
HAP1wt
ΔSLC30A1_1777_10
z-VAD(25µM)
VSV (MOI2)
Cleaved Caspase 3
Cleaved Caspase 7
GAPDH
75
15
15
e
HAP1
MOI 0,001
MOI 0,005
VSV-Δ51-GFP infected cells (%)
100
80
60
40
20
0
HAP1wt
ΔSLC30A1_1777_10

Suppl Fig 4

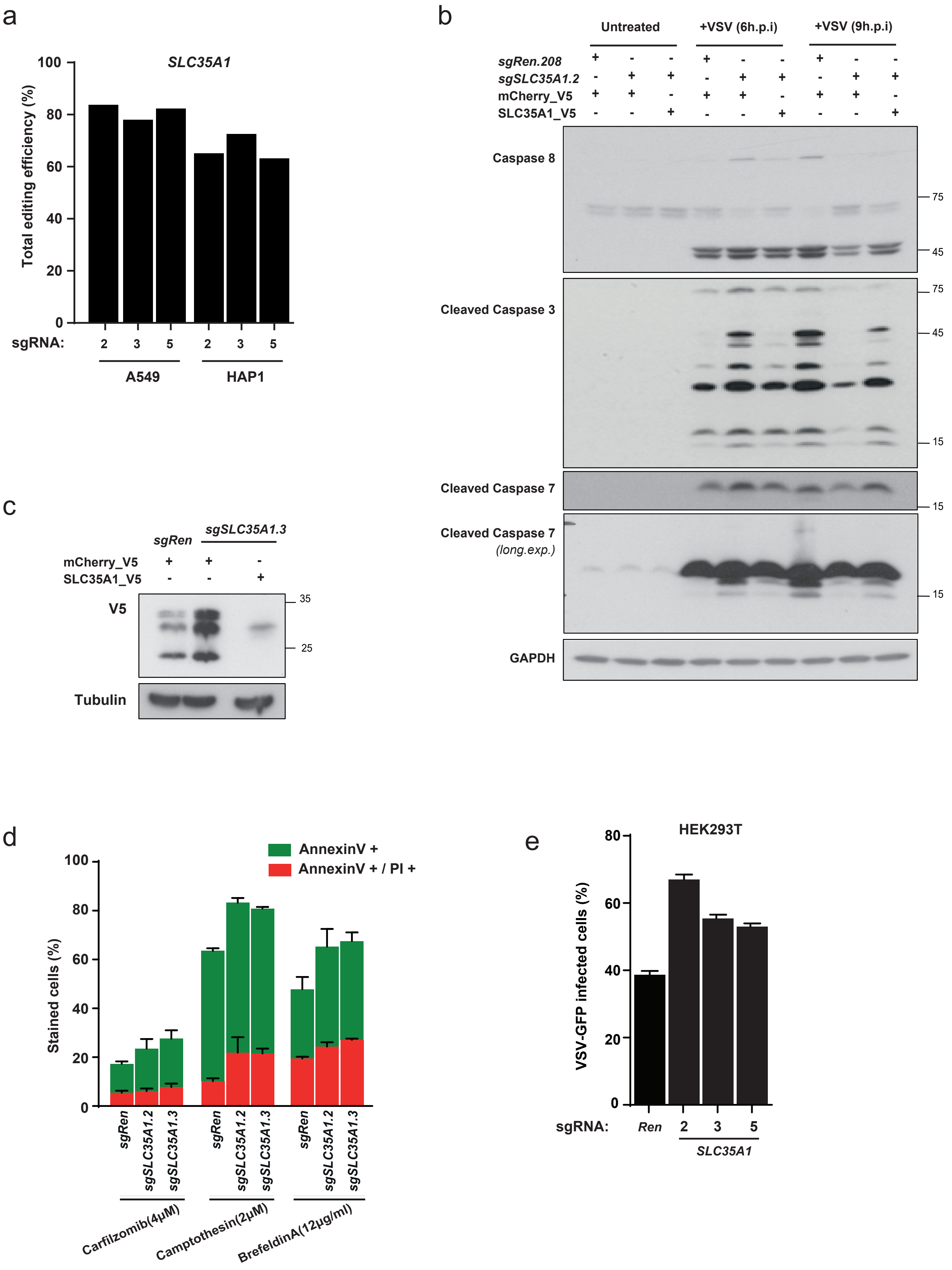
a
SLC35A1
Total editing efficiency (%)
100
80
60
40
20
0
sgRNA:
2 3 5 2 3 5
A549
HAP1
b
Untreated
+VSV (6h.p.i)
+VSV (9h.p.i)
sgRen.208
sgSLC35A1.2
mCherry_V5
SLC35A1_V5
Caspase 8
Cleaved Caspase 3
Cleaved Caspase 7
Cleaved Caspase 7 (long.exp.)
GAPDH
75
45
15
c
sgRen
sgSLC35A1.3
mCherry_V5
SLC35A1_V5
V5
Tubulin
35
25
d
AnnexinV +
AnnexinV + / PI +
Stained cells (%)
sgRen
sgSLC35A1.2
sgSLC35A1.3
Carfilzomib(4μM)
Camptothesin(2μM)
BrefeldinA(12μg/ml)
e
HEK293T
VSV-GFP infected cells (%)
sgRNA:
Ren
2
3
5
SLC35A1

### Full lenght immunoblots from Fig 2b

**Caspase 9**

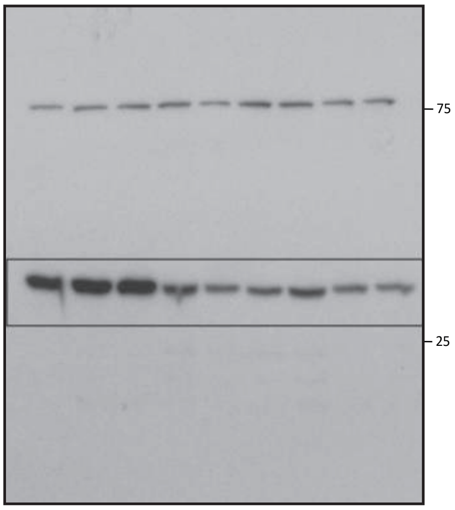


**Caspase 9**
*(long. exp.)*

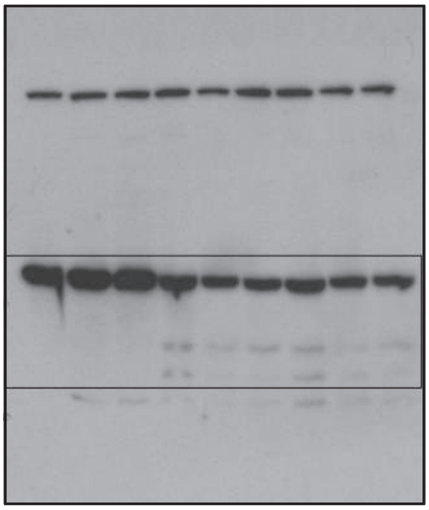

**Cleaved Caspase 7**

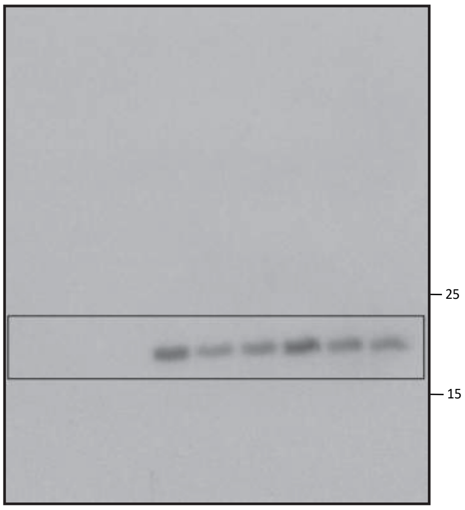


**Cleaved Caspase 3**

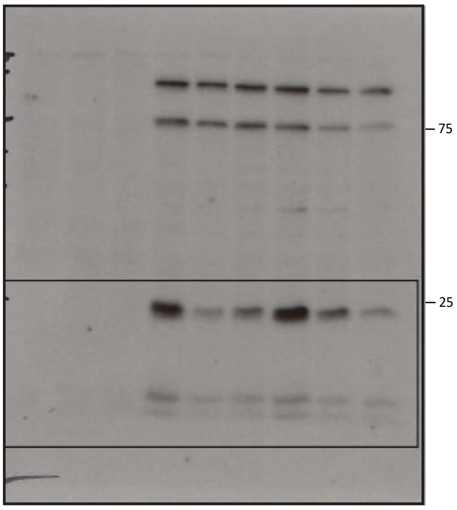


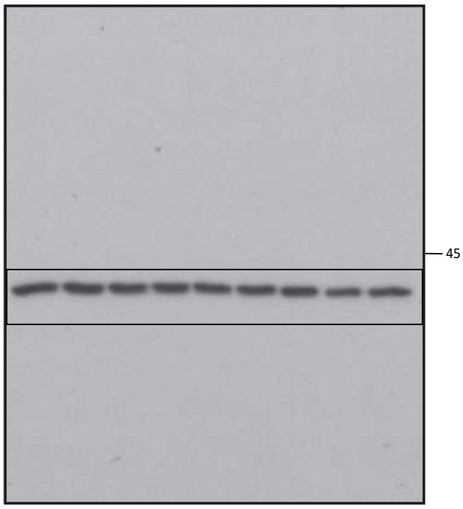


### Full lenght immunoblots from Fig 2c

**Caspase 8**

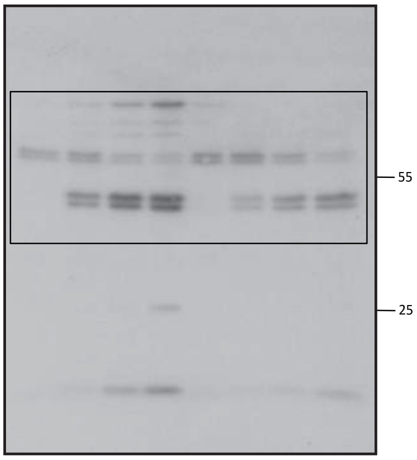


**Cleaved Caspase 7**

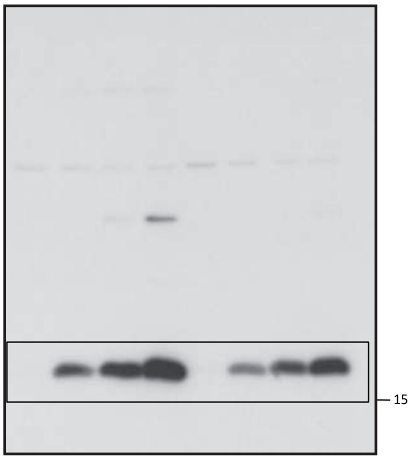


**Caspase 9**

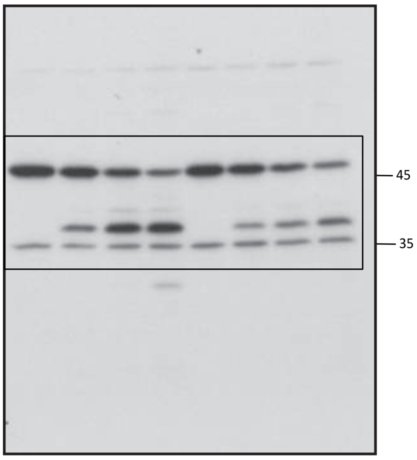

**Cleaved Caspase 3**

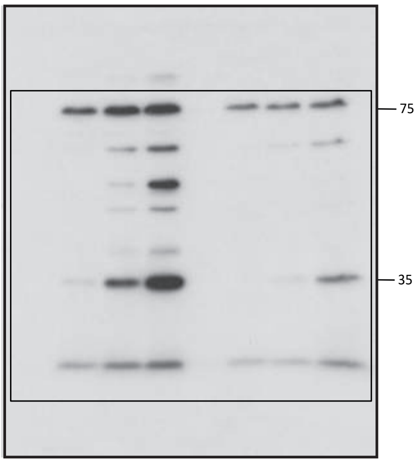


**GAPDH**

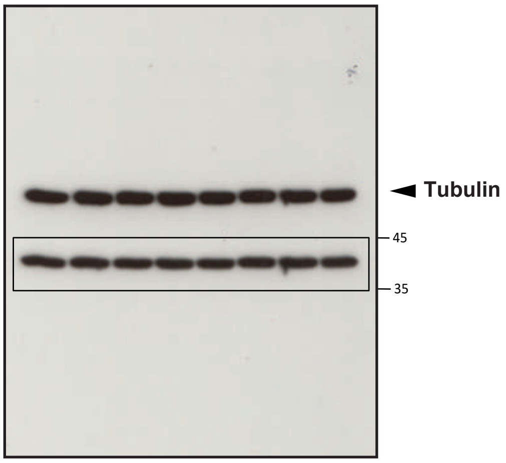


### Full lenght immunoblots from Fig 3d

**p-IRF3**

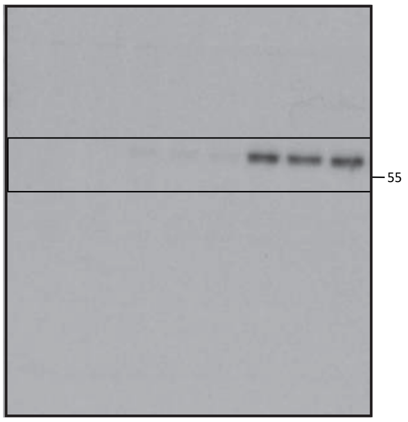


**p-IRF3**
*(long.exp.)*

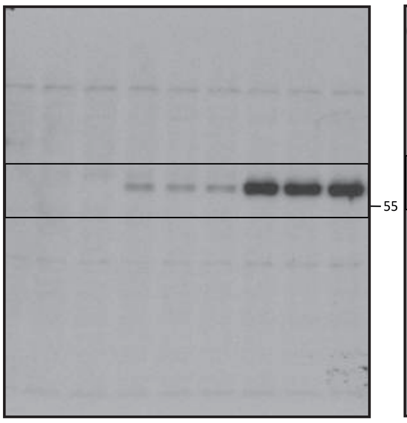


**IRF3**

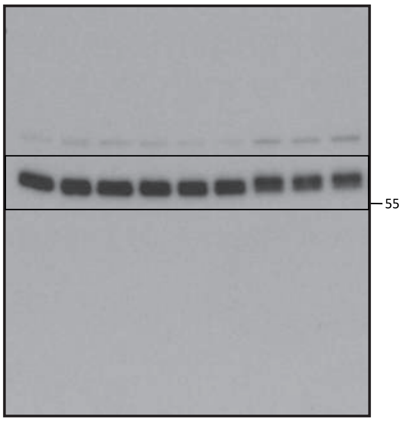


**p-STAT1**

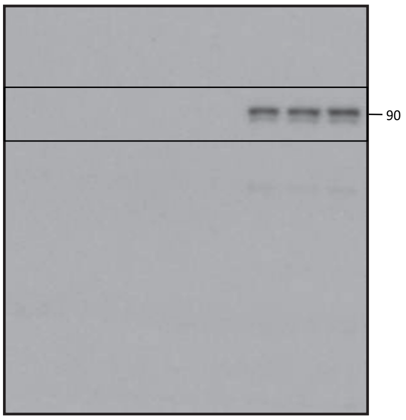


**p-STAT1**
*(long.exp.)*

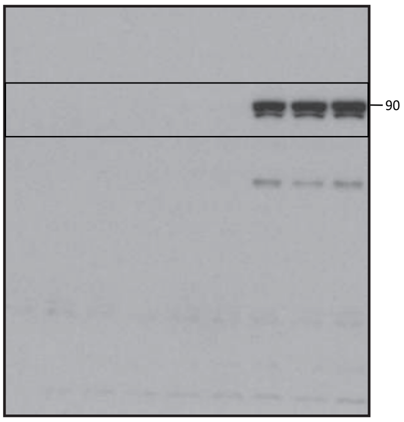


**STAT1**

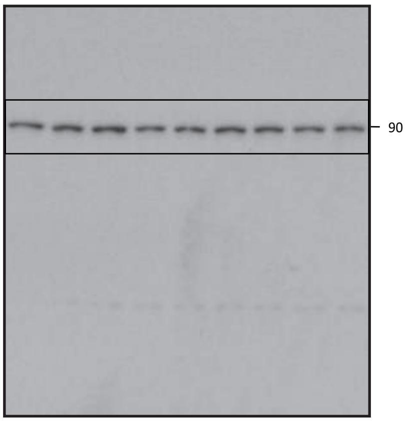


**p-p65**

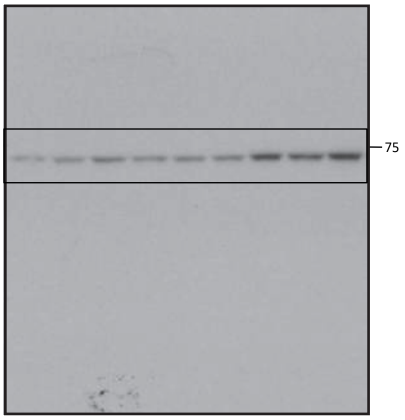


**p65**

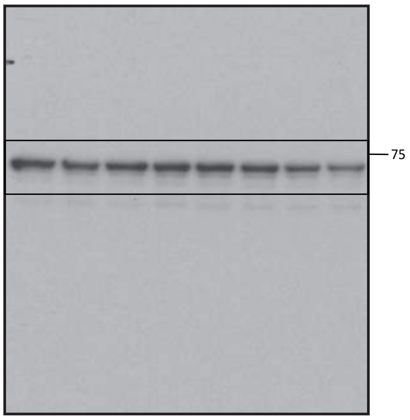


**GAPDH**

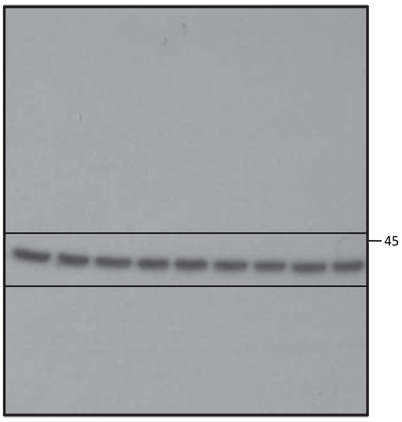

### Full lenght immunoblots from Fig 3f

**VSV_G**

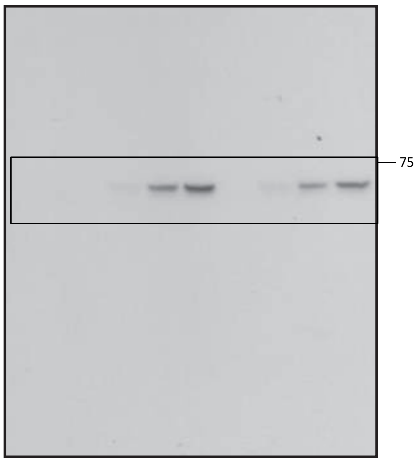


**GAPDH**

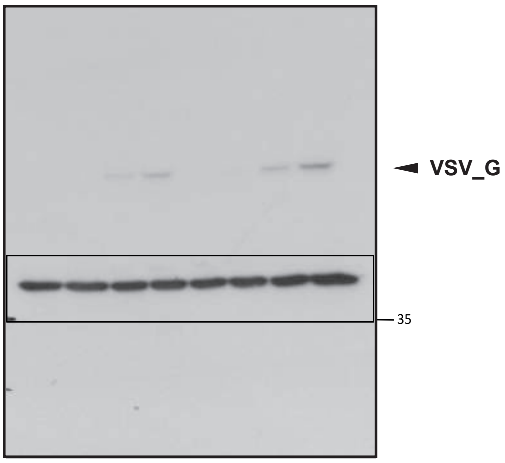


### Full lenght immunoblots from Fig 4b

**p-MEK**

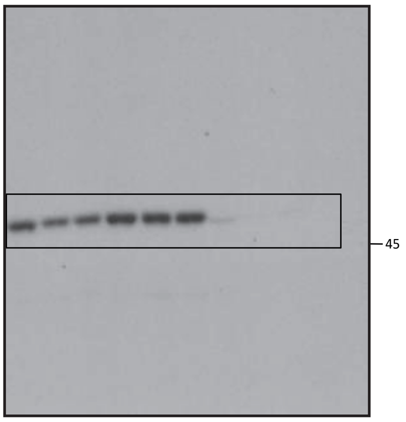


**p-MEK**
*(long.exp)*

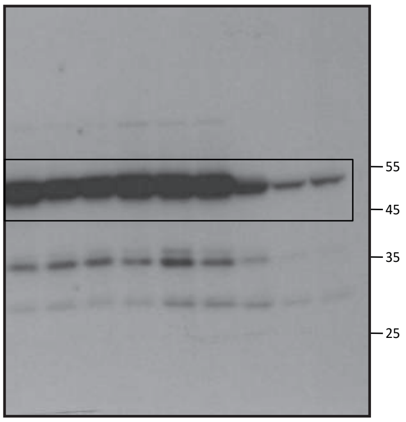


**MEK**

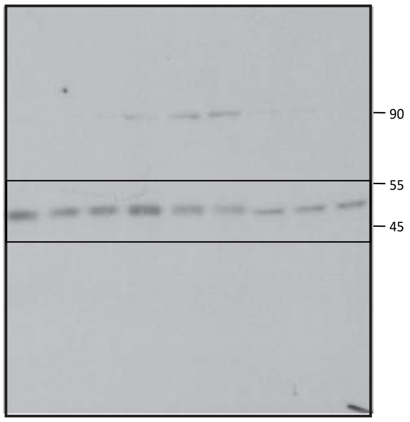


**p-ERK**

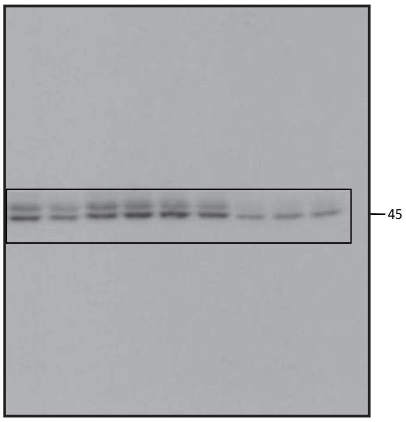


**ERK**

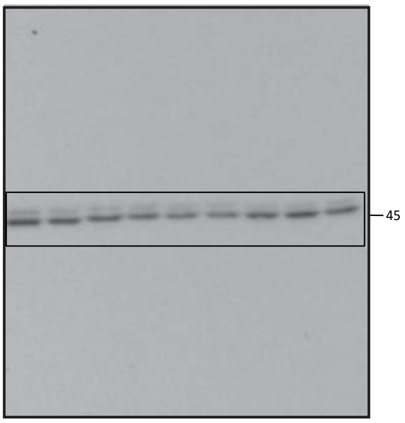


**ARF4**

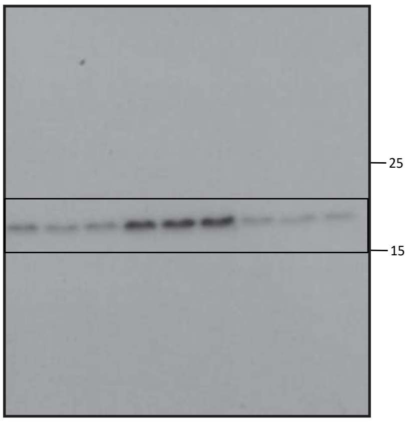


**GRP78**

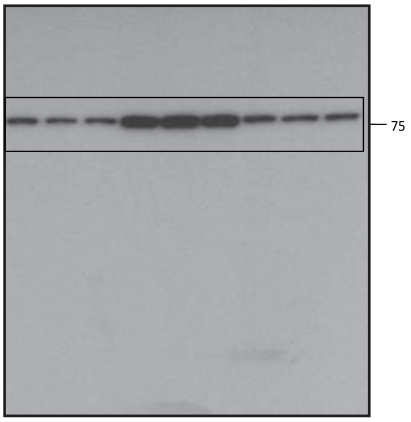


**CHOP**

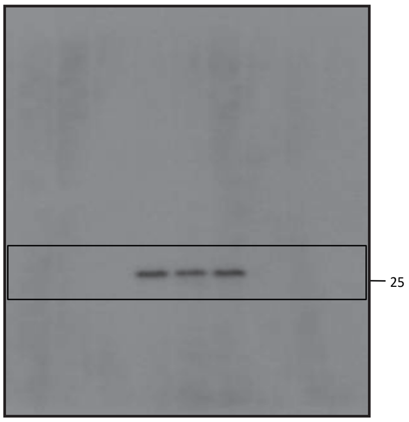


**GAPDH**

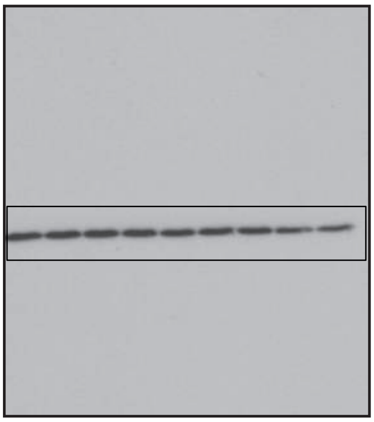

### Full lenght immunoblots from Suppl Fig 2c

**V5**

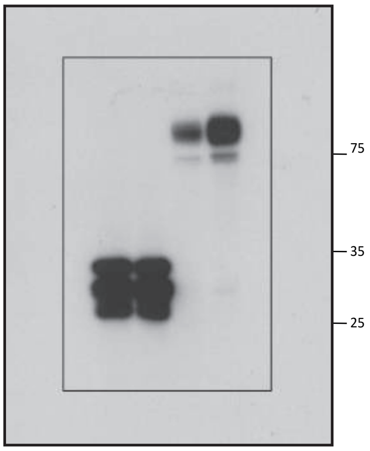


**GAPDH**

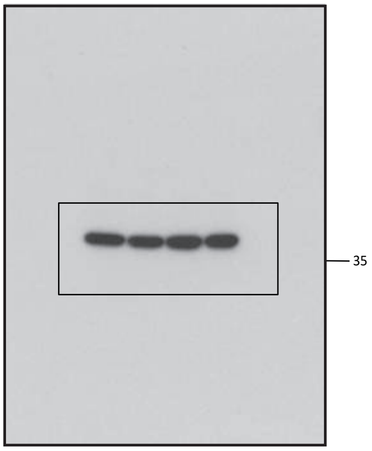


### Full lenght immunoblots from Suppl Fig 3d

**Cleaved Caspase 3**

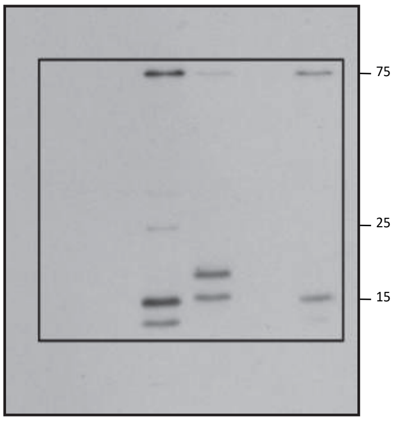


**Cleaved Caspase 7**

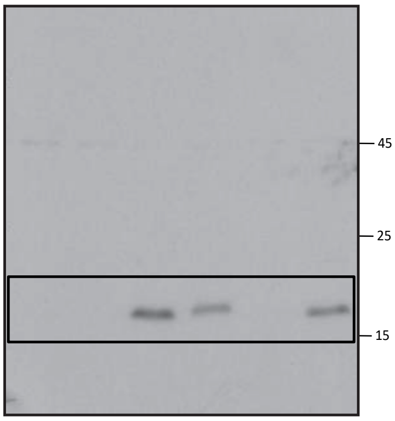


**GAPDH**

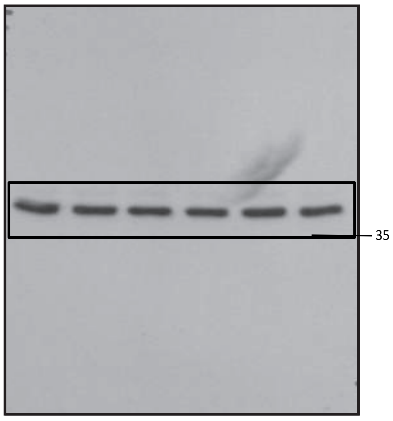


### Full lenght immunoblots from Suppl Fig 4b

**Caspase 8**

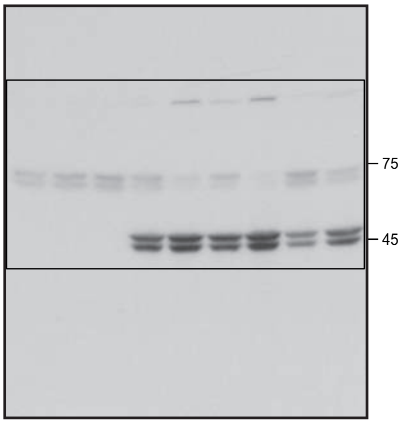


**Cleaved Caspase 3**

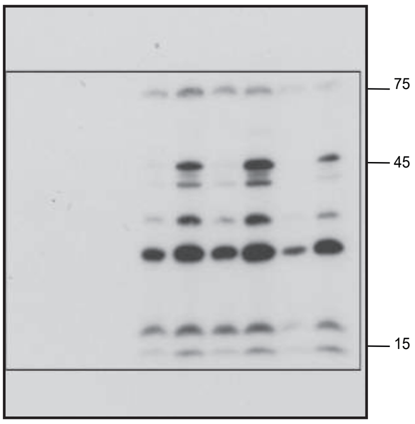


**Cleaved Caspase 7**

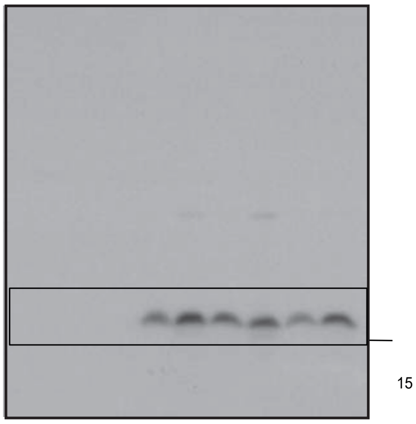


**Cleaved Caspase 7**
*(long.exp.)*

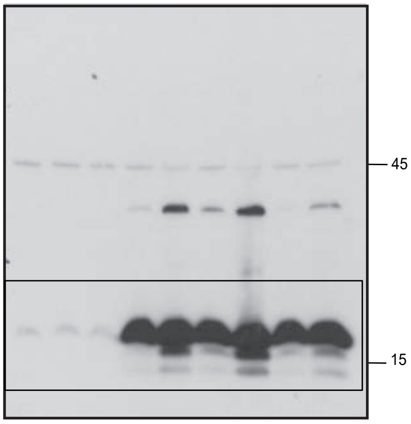


**GAPDH**

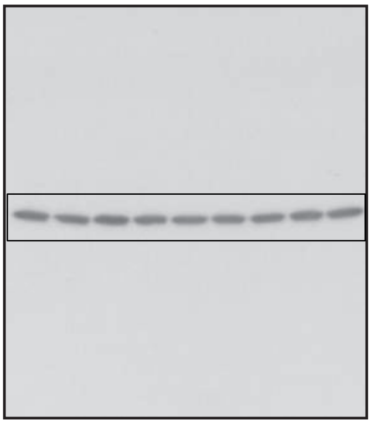

**Full lenght immunoblots from Suppl Fig 4c**

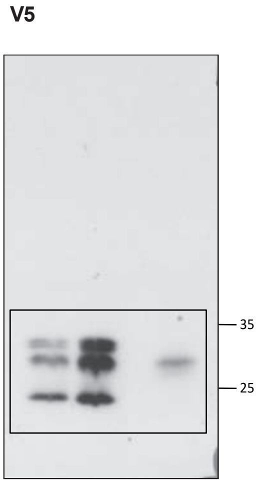


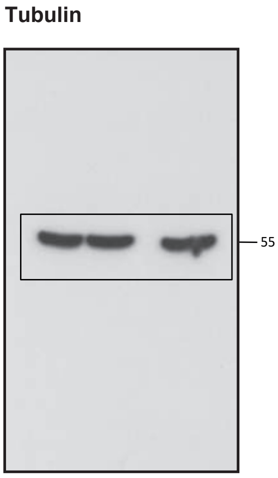
